# Implantation technique, safety and complications of robot-assisted stereoelectroencephalography exploration of the limbic thalamus in human focal epilepsy

**DOI:** 10.1101/2020.01.27.922195

**Authors:** Ganne Chaitanya, Andrew K. Romeo, Adeel Ilyas, Auriana Irannejad, Emilia Toth, Galal Elsayed, J. Nicole Bentley, Kristen O. Riley, Sandipan Pati

## Abstract

**Introduction:** Despite numerous imaging studies highlighting the importance of thalamus in surgical prognosis, human electrophysiological studies involving the limbic thalamic nuclei are limited. The objective of this study was to evaluate the safety and accuracy of robot-assisted stereotactic electrode placement in the limbic thalamic nuclei in patients with suspected temporal lobe epilepsy (TLE).

**Methods:** After obtaining informed consent, 24 adults with drug-resistant suspected TLE undergoing Stereo-EEG evaluation were enrolled in this prospective study. The trajectory of one electrode planned for clinical sampling the operculo-insular cortex was modified to extend to the thalamus, thereby preventing the need for additional electrode placement for research. The anterior thalamus (ANT) (N=13) and the medial group of thalamic nuclei (MED) (N=11), including mediodorsal (MD) and centromedian (CeM) were targeted. The post-implantation CT was co-registered to the pre-operative MRI, and Morel’s thalamic atlas was used to confirm the accuracy of implantation.

**Results:** Ten out of 13 (77%) in the ANT group and 10 out of 11 patients (90%) in the medial group had electrodes accurately placed in the thalamic nuclei. None of the patients had a thalamic hemorrhage. However, trace asymptomatic hemorrhages at the cortical level entry site were noted in 20.8% of patients and they did not require additional surgical intervention. SEEG data from all the patients were interpretable and analyzable. The trajectories for the ANT implant differed slightly from the medial group at the entry point i.e., precentral gyrus in the former and postcentral gyrus in the latter.

**Conclusions:** Using judiciously planned robot-assisted SEEG, we demonstrate the safety of electrophysiological sampling from various thalamic nuclei for research recordings, presenting a technique that avoids implanting additional depth electrodes, or comprising clinical care. With these results, we propose that if patients are fully informed of the risks involved, there are potential benefits of gaining mechanistic insights to seizure genesis, which may help to develop neuromodulation therapies.

## Introduction

Accurate localization of the epileptogenic zone (EZ) in temporal lobe epilepsy (TLE) is paramount to optimizing outcomes following surgical resection or ablation^44^. However, despite significant technological advancements in imaging and surgical tools, seizure-freedom outcomes have plateaued at 45-65% over the last decade, necessitating a continued investigation of how distributed brain regions interact to cause seizures^39^. Growing evidence suggests that networks of functionally and structurally connected areas, both within and outside of mesial temporal structures, contribute to EZ’s^4,10,19,32^. Among extra-temporal structures implicated in seizure inception, increased thalamo-temporal structural and functional connectivity independently predicts poor surgical outcomes^17^. Experimental studies in preclinical models of limbic epilepsy support the hypothesis that thalamus can regulate limbic seizures, and the stage at which ictogenesis is regulated (i.e., initiation, propagation, or termination) depends on the functional connectivity of the thalamic nuclei with limbic structures^6,9,36^. Furthermore, modulation of the “limbic” thalamic nuclei (i.e. the anterior nucleus -ANT, midline and mediodorsal -MD, and the intralaminar centromedian -CeM) through chemical, optogenetic, or electrical perturbation can interrupt focal seizures^6,16,25,37^. However, despite numerous preclinical and imaging studies highlighting the importance of thalamus in surgical prognosis^17^, there are very limited human electrophysiological studies targeting the limbic thalamus.

Over the last decade, clinicians have implanted subdural grids, hybrid macro-micro depth electrodes, and laminar electrodes for high-density intracranial recordings from both the superficial and deep cortex^40,43^. Although the thalamus is likely a rich source of data about seizure regulation, propagation, and sleep dysfunction^12,15^, progress in this area is relatively slow due to ethical and safety concerns. However, future therapeutic developments may be highly informed by understanding thalamo-temporal causal interactions during seizures. Here, we present the technical nuances, safety, and accuracy data from stereoelectroencephalography (SEEG) implantation of electrodes into the limbic thalamus during the presurgical evaluation of patients with suspected TLE.

## Methods

### Patient selection

Enrolled patients included those deemed eligible for SEEG after consensus recommendation from a multi-disciplinary epilepsy conference consisting of neurologists, neurosurgeons, neuropsychologists, and nurses. Adults with drug-resistant suspected TLE undergoing SEEG evaluation were eligible to participate in the study. Surgeons modified the trajectory of one electrode planned for clinical sampling to extend to the thalamus, obviating the need to implant an additional electrode for thalamic sampling. A five-stage evaluation process helped streamline the process of recruiting considerably homogenous groups of patients with suspected TLE (mesial and/or temporal plus) who would receive the thalamic implant (Fig 1). The investigators approached the potential candidates for thalamic SEEG at the outpatient follow-up clinic visit, and written informed consent was obtained in accordance with protocols approved by the University of Alabama Birmingham Institutional Review Board.

**Figure 1:**
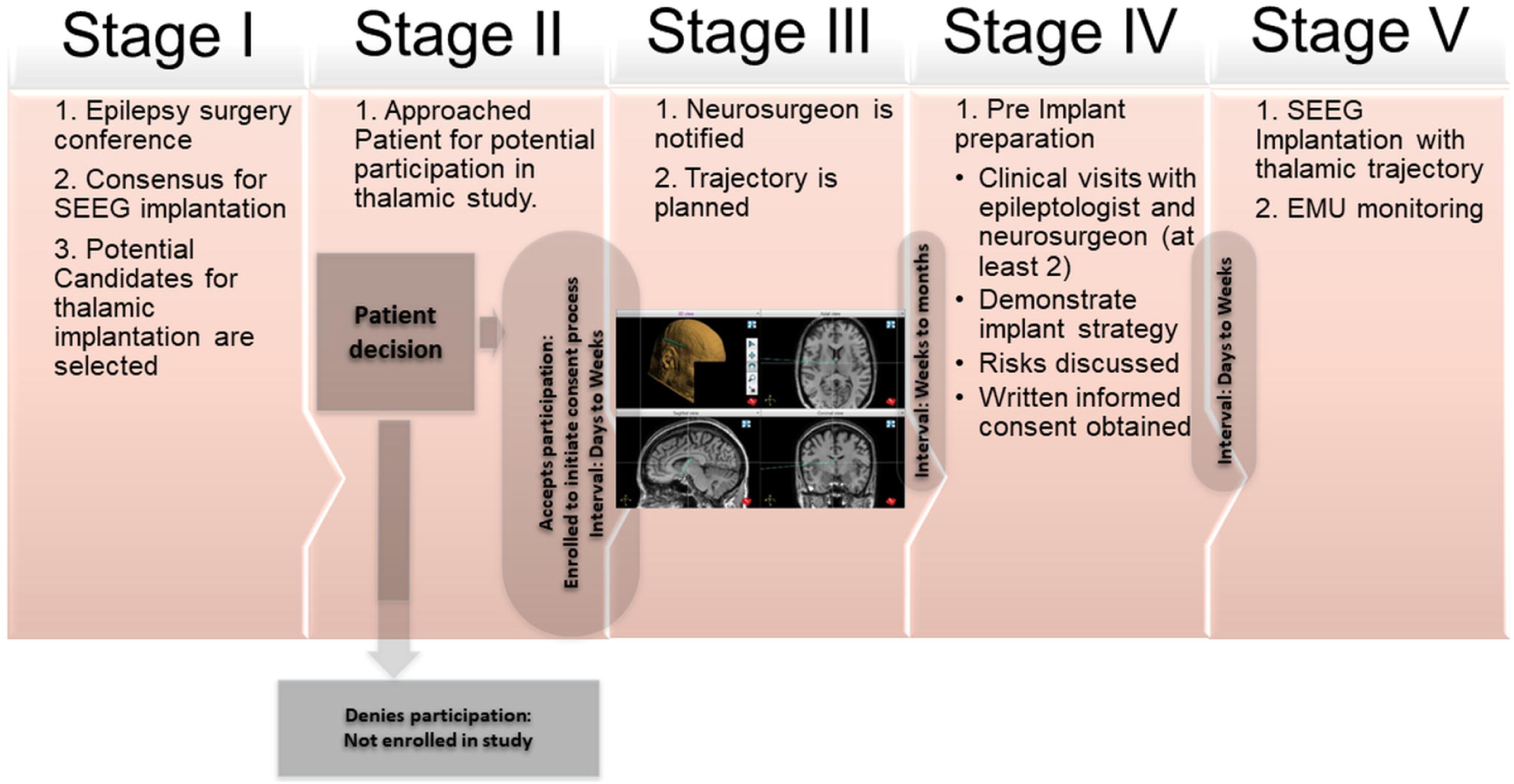
Patient selection and SEEG implantation: A multi-stage process to obtain informed consent from eligible patients scheduled to undergo SEEG investigation for localization of epilepsy

### Thalamic trajectory

In this study, the limbic thalamic nuclei that were targeted were: 1) anterior thalamus (ANT) including anterior ventral (AV), anterior dorsal (AD) and anterior medial subnuclei (AM); 2) medial thalamus (MED) including mediodorsal (MD) and centromedian subnuclei (CeM). Neurosurgeons planned trajectories using T1-weighted sequences with gadolinium contrast magnetic resonance imaging (MRI) using robotic stereotactic planning software (ROSA^©^, Medtech, Montpellier, France). MRI scans were acquired using epilepsy protocol in Philips Achieva scanner 1.5T (matrix size: 384× 384mm/ 432×432mm, SliceThickness: 1.2mm, TR: 7ms, TE: 3ms, interslice gap:1.2mm, FlipAngle: 8), Philips Achieva scanner 3T (matrix size: 256×256/528×528, SliceThickness: 1mm/1.2mm, TR: 6ms, TE: 3ms/4ms, interslice gap:1mm/1.2mm, FlipAngle: 8/9), Philips Ingenia scanner 1.5T(matrix size: 512× 512mm/ 432×432mm, SliceThickness: 1mm, TR: 7ms, TE: 3ms, interslice gap:1mm, FlipAngle: 8), Philips Ingenia scanner 3T(matrix size: 528× 528mm/ 432×432mm, SliceThickness: 1.2mm, TR: 7ms, TE: 3ms, interslice gap:1.2mm, FlipAngle: 9), SIEMENS Skyra scanner 3T(matrix size: 256×256mm, SliceThickness: 0.9mm, TR: 7ms, TE: 3ms, interslice gap:0mm, FlipAngle: 8). Post-explant CT scans were also obtained to look for postoperative hemorrhage or edema. Post-operative hemorrhage if found was graded using McGovern’s SEEG hemorrhage grading^23^. Based on a visual reference to the Morel’s thalamic atlas, the nuclei were determined to be located in relation to the following landmarks^21^ while planning the trajectory. The ANT was identified by its close relationship to the foramen of Monro and the venous angle formed by the confluence of thalamostriate and septal veins lying immediately posterior and lateral (Fig 2). Extending a trajectory incorporating the frontal operculum and insula, the thalamic region of interest was targetedwithout requiring an additional electrode. For the medial group of thalamic nuclei, neurosurgeons targeted the ventral midline thalamus near the massa intermedia (MI), providing recordings from the central-medial or mediodorsal nuclei. The broadest midline nuclear segment was anterior and ventral to the MI, where the reuniens and central median nuclei are located^33^. The MI could be visualized in five of the eight medial thalamic groups of patients. When not distinct, the anteroposterior mid-point at the level of AC-PC plane was chosen ^5^. The inferior and nearly parallel relationship with the ANT provided further anatomic confirmation.

**Figure 2:**
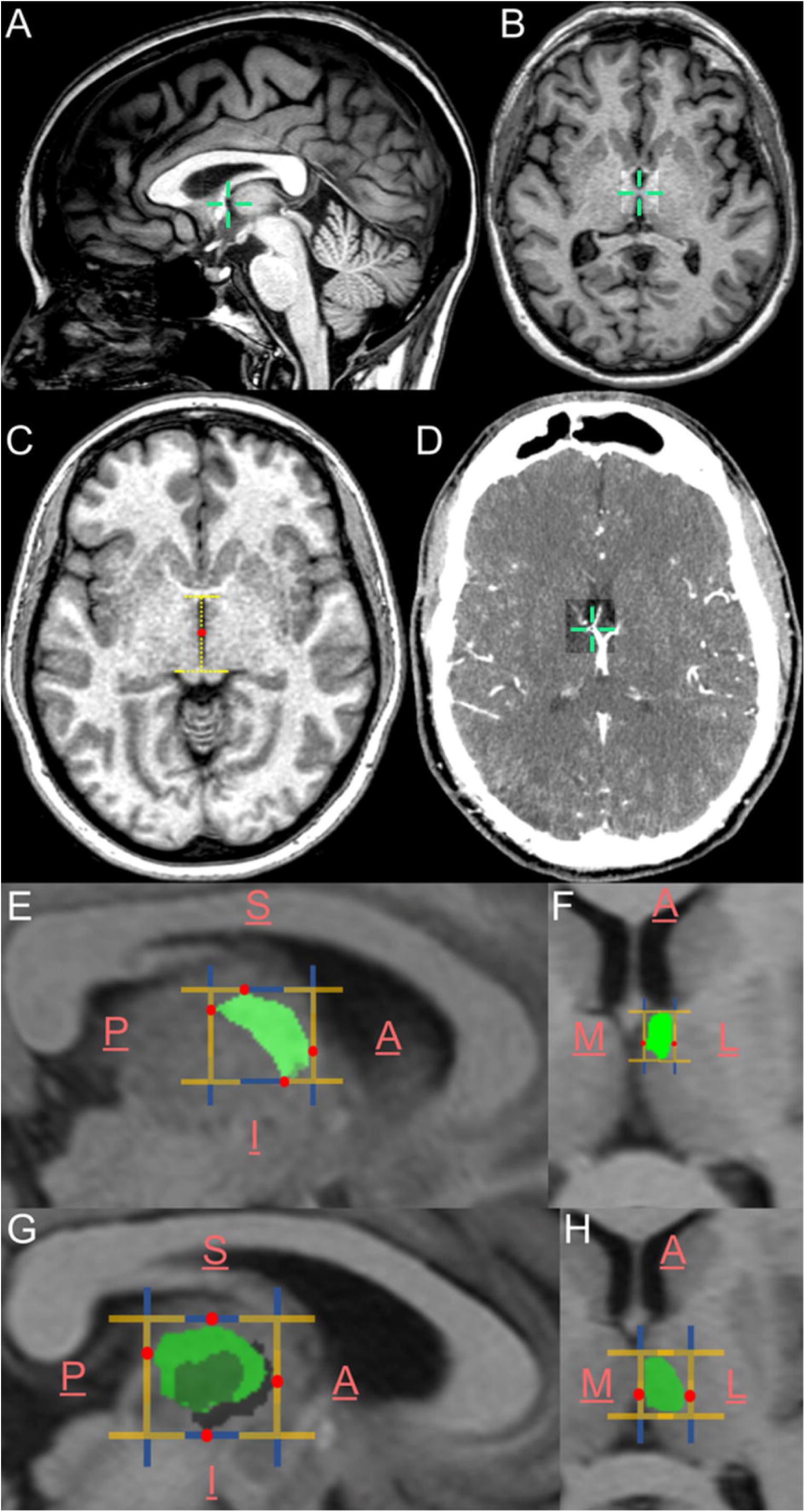
Landmarks for targeting the thalamic nuclei: (A) Foramen of Munroe (FM) as the antero-medial extent, (B) the massa intermedia (MI) as the medial extent, (C) the midpoint of the anntero-posterior length of the third ventricle in axial view as the inferior extent (measured at the level of superior colliculus and the venous angle as the antero-medial extent (confluence of anterior septal vein (ASV), thalamostriate vein (TSV), and internal cerebral vein (ICV) were the major landmarks used while targeting the electrodes. (E&F) represent the probable extents of ANT as per Morel’s atlas. (G&H) represent the probable extents of medial nuclear group (CeM+MD) as per Morel’s atlas. (Anterior-A, posterior-P, medial-M, lateral-L, superior-S and inferior-I).

Following SEEG implantation, patients were initially monitored in the neurosurgical intensive care unit for 24 hours. A high-resolution post-implantation head CT was obtained within 24 hours. Subsequently, the patients were transferred to the epilepsy monitoring unit for seizure localization and mapping.

### Measuring Accuracy

Post-implantation CT scans were acquired using Philips Brilliance64 and Brilliance16P scanners with matrix size: 512×512, SliceThickness: 1mm, KVP: 120, interslicegap: 1mm, ExposureTime: 1550s/727s. The CT scnas were co-registered to the pre-operative MRI using Advanced Normalization Tools (ANTs) (Fig 3)^3^. Electrodes were localized using Lead-DBS software (www.lead-dbs.org), and trajectories were mapped using iElectrodes software^7,18^. Co-registered images were normalized to ICBM152-2009b Nonlinear Asymmetric using nonlinear diffeomorphic normalization algorithms, and brain shift corrections were performed using Schönecker normalizations providing refined registration of subcortical structures^35^. Registrations were checked manually for errors in 3DSlicer. The data were then visualized using Morel’s atlas^21^. To map the trajectories, the co-registered and normalized CT and MRI scans were imported along with the corresponding CT mask into iElectrodes and registered with the AAL2 atlas to identify their cortical-subcortical locations^28,41^. As a final step in establishing the accuracy of the implant strategy from our experience, we estimated post-implant accurate location of the thalamic target along with the linear component distances along the X, Y, and Z axes and the Euclidean distances from the landmarks.

**Figure 3:**
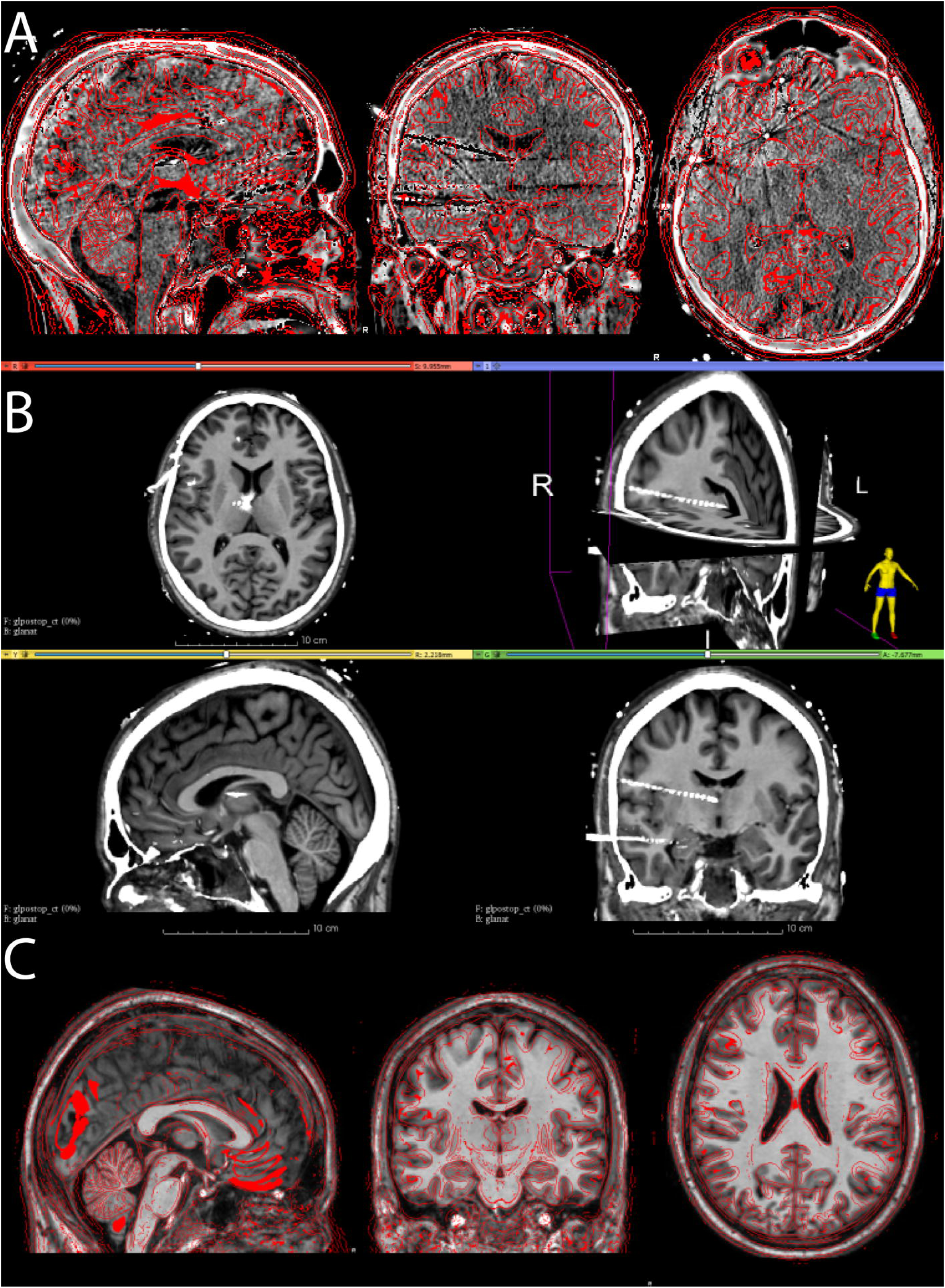
Co-registration strategy: (A) The post-implantation CT scan was coregistered onto the pre-operative MRI to determine postoperative target of the thalamic targets using Advanced Normalization Tools (ANTs) in Lead-DBS. (B) Following coregistration a manual verification of the anatomical landmarks and the overlap of the inner and outer tables of MRI and CT are used to ensure accurate coregistration. In particular, the most probable location of the thalamic electrode is confirmed.(C) The coregistered MRI and CT scan were then normalized to the MNI space. The coregistered images were then resolved for finer registration and brainshifts using Schönecker normalization algorithm to improve the registration of subcortical structure (3D Slicer).

## Results

### Patient Demographics

Details regarding patient demographics are shown in **Table 1**. A total of 24 patients underwent thalamic SEEG implantation. The ANT was targeted in 13 patients (∼54%), and the medial group nuclei were targeted in 11 patients (∼46%) (M:F=11:13, mean age at implantation= 40±11years). MRI brain was negative for any epileptogenic lesion in 10 patients (41.6%), while 4 patients (16.7%) had hippocampal sclerosis, 5 (20.8%) had hippocampal sclerosis with additional extra-hippocampal pathology, and the remaining 5 had extrahippocampal pathology only (20.8%).

**Table 1:**
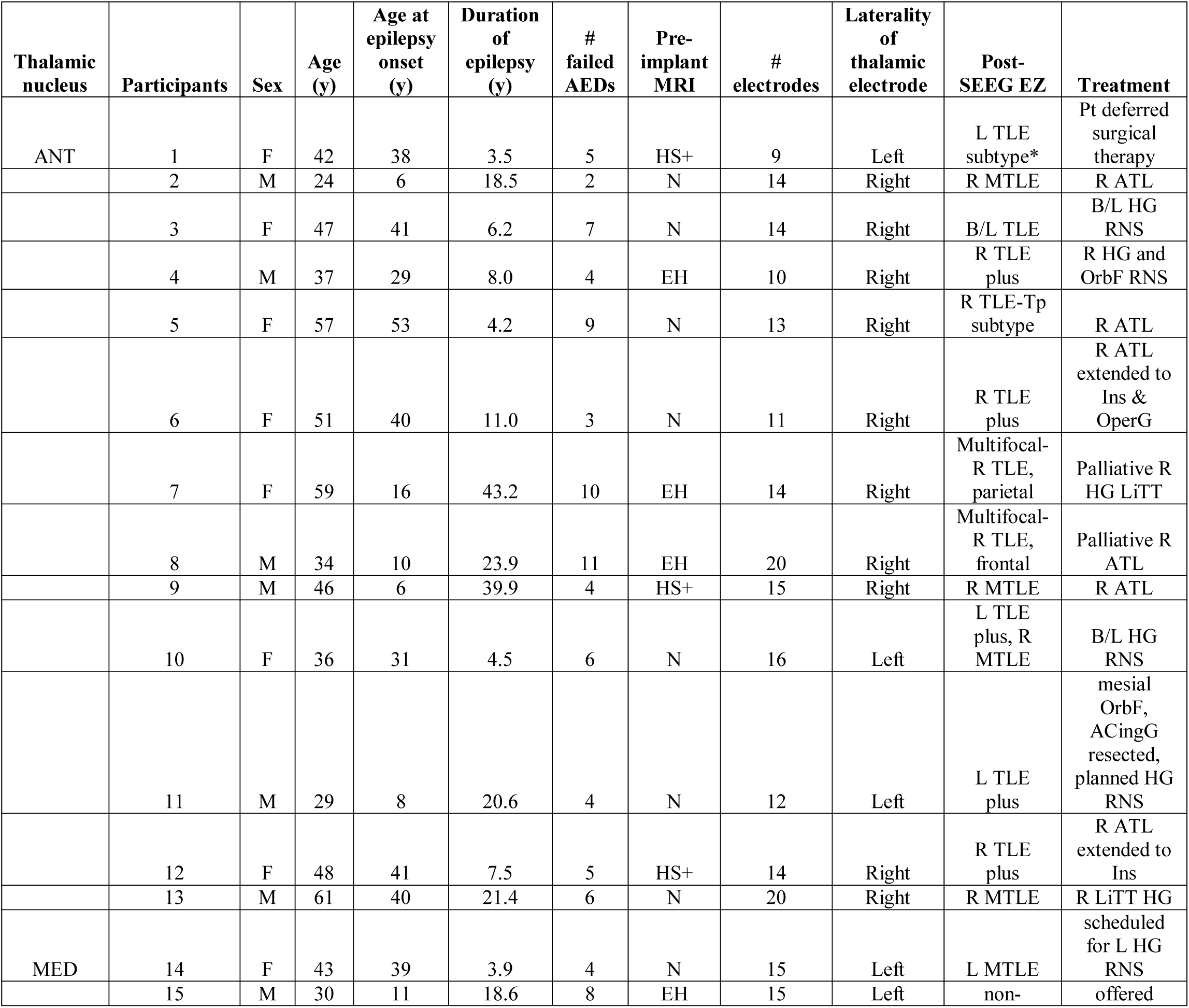
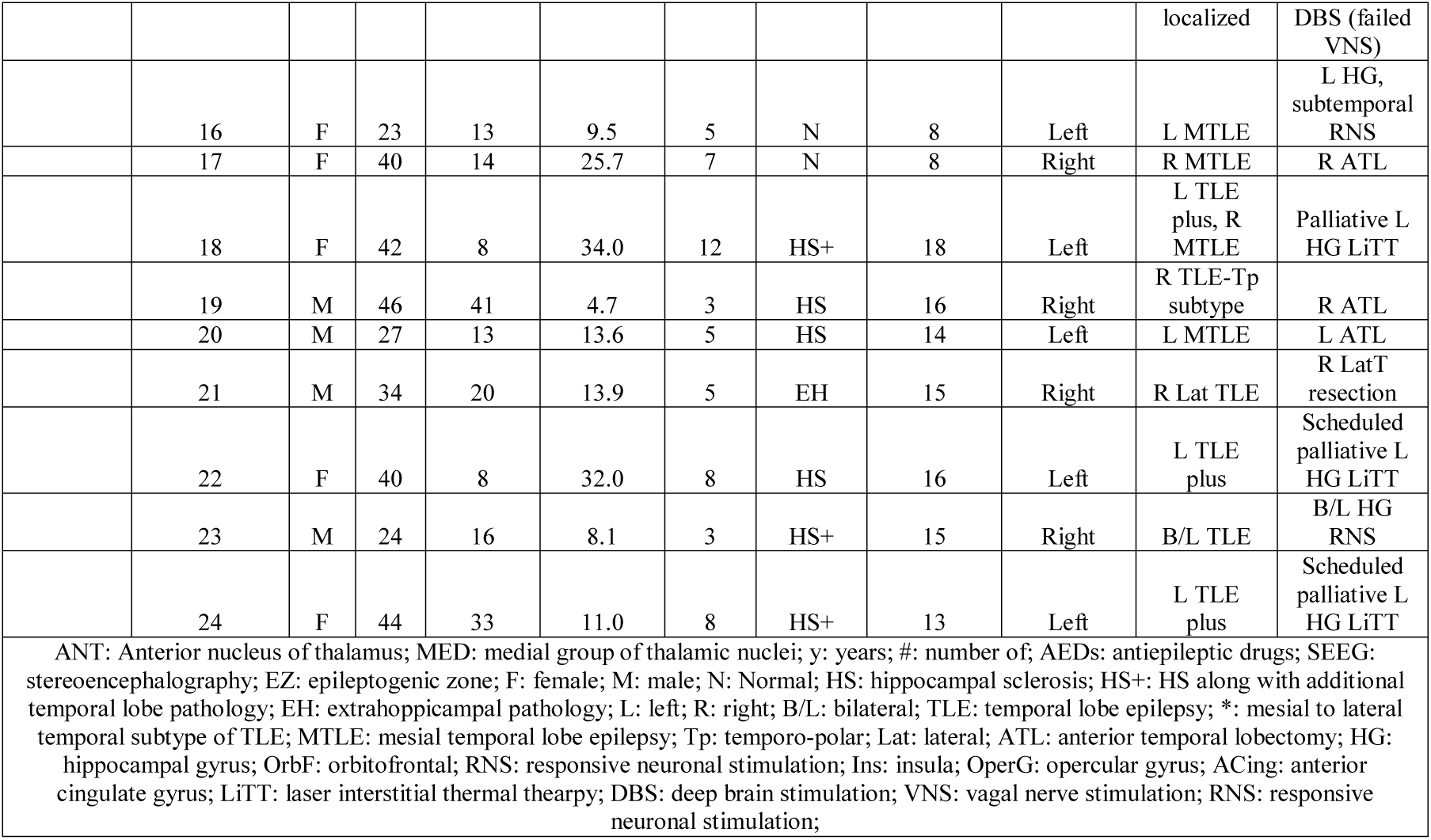
Patient demographics: The clinical presentation of the 24 participants are enumerated along with the post-SEEG localization of the seizure pathology and the treatment strategy.

### SEEG Implantation and Complications

On an average, each patient received a median of 172 contacts (range = 102-274). SEEG implantation and outcome data are presented in **Table 1**. None of the patients in our study group had thalamic hemorrhage or edema. Asymptomatic subarachnoid (SAH), subdural (SDH) and intracranial hemorrhages (ICH) were noted close to the entry site of the electrodes in 5/24 (21%) patients (Fig 4-B). All were grade 1-2 hemorrhages as per McGovern’s SEEG hemorrhage grading^23^ (consisting of a small intracranial bleed, either close to or away from eloquent cortex), which were low-grade hemorrhages with a lesser probability of being symptomatic. On evaluating the three major risk factors as identified in study by McGovern et al., we found no significant difference in gender, age or number of electrode implants for patients with and without hemorrhage [(M:F=4:1, chi-square=3.8, p=0.051); (Age: hemorrhage group: 42.7±7years, No hemorrhage group: 39.4±11years, t=0.58, p=0.56); (Number of electrodes: hemorrhage group: 14±4 electrodes, No hemorrhage group: 14±3 electrodes, t=0.34, p=0.73)]. Asymptomatic vasogenic edema in temporal or parietal lobe was noted in 3/24 patients. Overall, 34% of patients had asymptomatic hemorrhage or edema in the post-explant CT scans. Follow-up CT scans showed resolution of these findings. None of the patients had any symptomatic hemorrhage or required any surgical interventions to treat hemorrhage.

**Figure 4:**
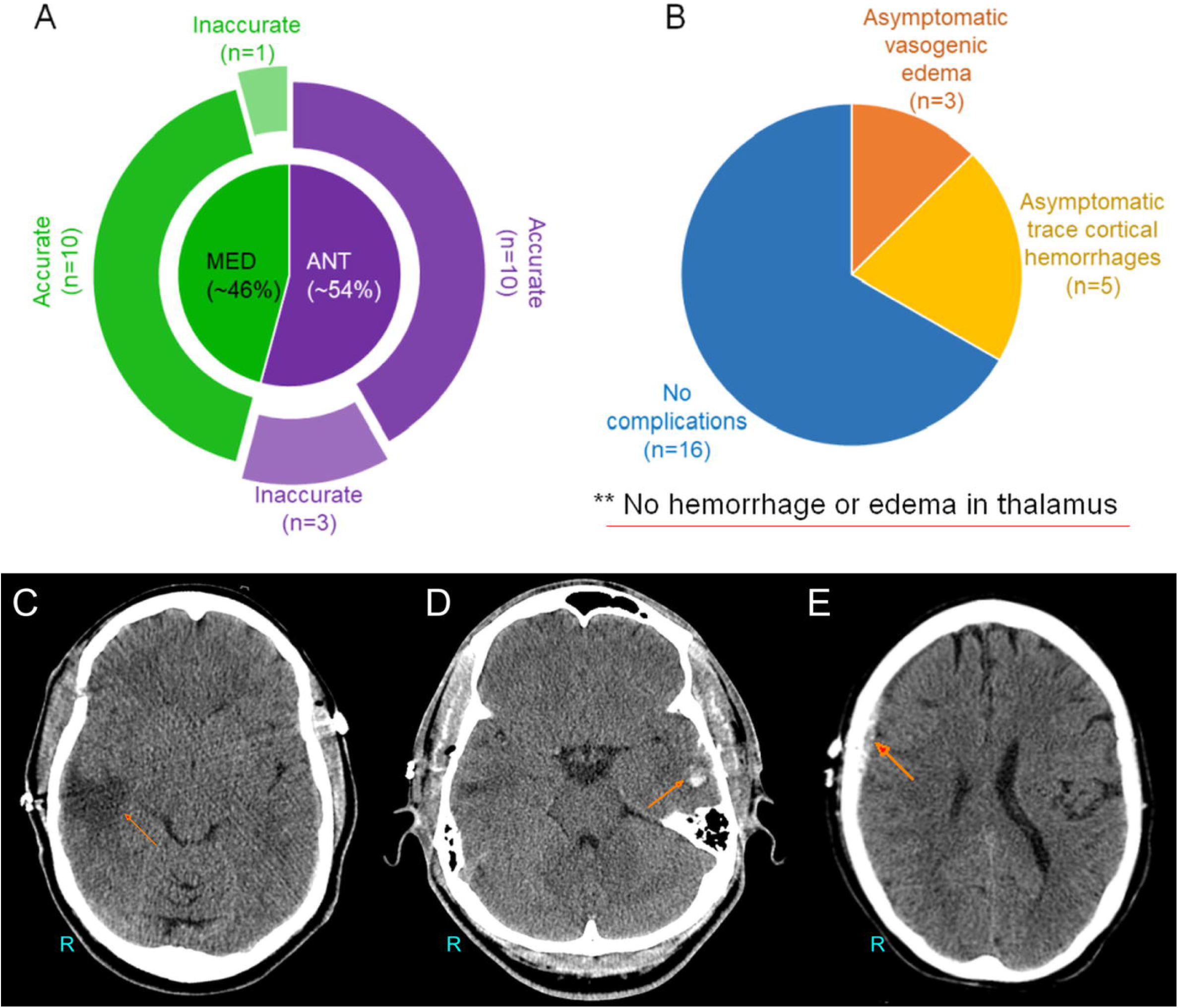
Thalamic implant and complication rates: (A) graph represents the number of patients receiving ANT or MED thalamic implants. (B) graph represents the number of patients who had post-operative complications in terms of asymptomatic vasogenic edema and asymptomatic hemporrhages at the proximity of the entry site of the electrodes. (C) Postexplant CT scan showing asymptomatic vasogenic edema in the right temporal lobe. (D) Postexplant CT scan showing asymptomatic left temporal intracranial hematomoa. (E) Postexplant CT scan showing asymptomatic right frontal extradural hematoma.

### Targeting Accuracy

Of the 13 patients who were planned for ANT implantation, 10 (77%) had confirmed localization in the ANT (Fig. 4-A). In one of the patients, the electrode passed through ipsilateral ANT and crossed the midline to the opposite side thalamus traversing through the MD nucleus. In another patient, the electrode target stopped short by 4mm and was situated in the ventral lateral nucleus of the thalamus. In the third patient the electrode target was situated 3mm anterior to the ANT in the anterior fornix.

For the medial group of nuclei (n=11), the massa intermedia is used for localization. Ten electrodes (90%) were situated in the medial group of nuclei (CeM and MD). In one patient, the electrode target stopped short by 8mm from the midline and was situated in the ventral medial thalamic nucleus.

Post-implant accuracies in X, Y, and Z axes and Euclidean errors are presented in **Table 2**. The trajectories for the ANT implant differed slightly from the medial group at the entry point that was in the precentral in the former and postcentral gyrus in the latter (Fig 5).

**Table 2:** Measurements of electrode targets to anatomical landmarks: The measurements (in mm) between the electrode target and the anatomical landmarks are provided in terms of actual 3D coordinates of the electrodes, their linear components vectorized on x,y and z axes and finally the 3D Euclidean distance in space between the target tip and the anatomical landmark.

| ANTERIOR THALAMUS | TARGET |  |  | FM |  |  | MI |  |  | TV |  |  |
| --- | --- | --- | --- | --- | --- | --- | --- | --- | --- | --- | --- | --- |
|  | X (mm) | Y (mm) | Z (mm) | X (mm) | Y (mm) | Z (mm) | X (mm) | Y (mm) | Z (mm) | X (mm) | Y (mm) | Z (mm) |
| LOCATION | M±SD | M±SD | M±SD | M±SD | M±SD | M±SD | M±SD | M±SD | M±SD | M±SD | M±SD | M±SD |
| Left (n=1) | -5 | -5 | 8 | -4 | 1 | 5 | 0 | -12 | 10 | 0 | -12 | -4 |
| Right (n=9) | 2±5 | -6±2 | 6±7 | 2±1 | 0±1 | 3±2 | 0±0 | -11±2 | 2±5 | 0±0 | -12±1 | -4.1±1 |
| Linear Component Distance | TARGET to FM |  |  | TAREGT to MI |  |  | TARGET to TV |  |  |  |  |  |
| Left | 1 | 6 | 3 | 5 | 7 | 2 | 5 | 7 | 12 |  |  |  |
| Right | 3±4 | 6±2 | 6±3 | 5±2 | 5±2 | 7±5 | 5±2 | 7±2 | 11±5 |  |  |  |
| Euclidian Distance (mm) | to FM | to MI | to TV |  |  |  |  |  |  |  |  |  |
| Left | 7 | 9 | 15 |  |  |  |  |  |  |  |  |  |
| Right | 10±3 | 11±3 | 15±2 |  |  |  |  |  |  |  |  |  |
| MEDIAL THALAMUS | TARGET |  |  | FM |  |  | MI |  |  | TV |  |  |
|  | X (mm) | Y (mm) | Z (mm) | X (mm) | Y (mm) | Z (mm) | X (mm) | Y (mm) | Z (mm) | X (mm) | Y (mm) | Z (mm) |
| LOCATION | M±SD | M±SD | M±SD | M±SD | M±SD | M±SD | M±SD | M±SD | M±SD | M±SD | M±SD | M±SD |
| Left (n=8) | -3±1 | -10±3 | 2±4 | -3±1 | -1±1 | 2±2 | 0±0 | -10±2 | 2±5 | 0±0 | -12±1 | -4±1 |
| Right (n=2) | 5±3 | -12±1 | 1±4 | 2±1 | 0±2 | 3±4 | 0±0 | -10±1 | -2±1 | 0±0 | -13±2 | -51 |
| Linear Component Distance | TARGET to FM |  |  | TAREGT to MI |  |  | TARGET to TV |  |  |  |  |  |
| Left | 0±1 | 9±3 | 4±2 | 2±1 | 2±3 | 2±3 | 2±1 | 3±3 | 7±3 |  |  |  |
| Right | 4±3 | 12±2 | 3±2 | 5±3 | 2±1 | 3±3 | 5±3 | 1±2 | 6±5 |  |  |  |
| Euclidian Distance (mm) | to FM | to MI | to TV |  |  |  |  |  |  |  |  |  |
| Left | 10±2 | 5±3 | 8±4 |  |  |  |  |  |  |  |  |  |
| Right | 13±2 | 7±2 | 9±3 |  |  |  |  |  |  |  |  |  |
| FM: Foramen of Munro; MI: massa intermedia; TV: antero-posterior mid-point of third ventricle; M: mean; SD: Standard deviation |  |  |  |  |  |  |  |  |  |  |  |  |

**Figure 5:**
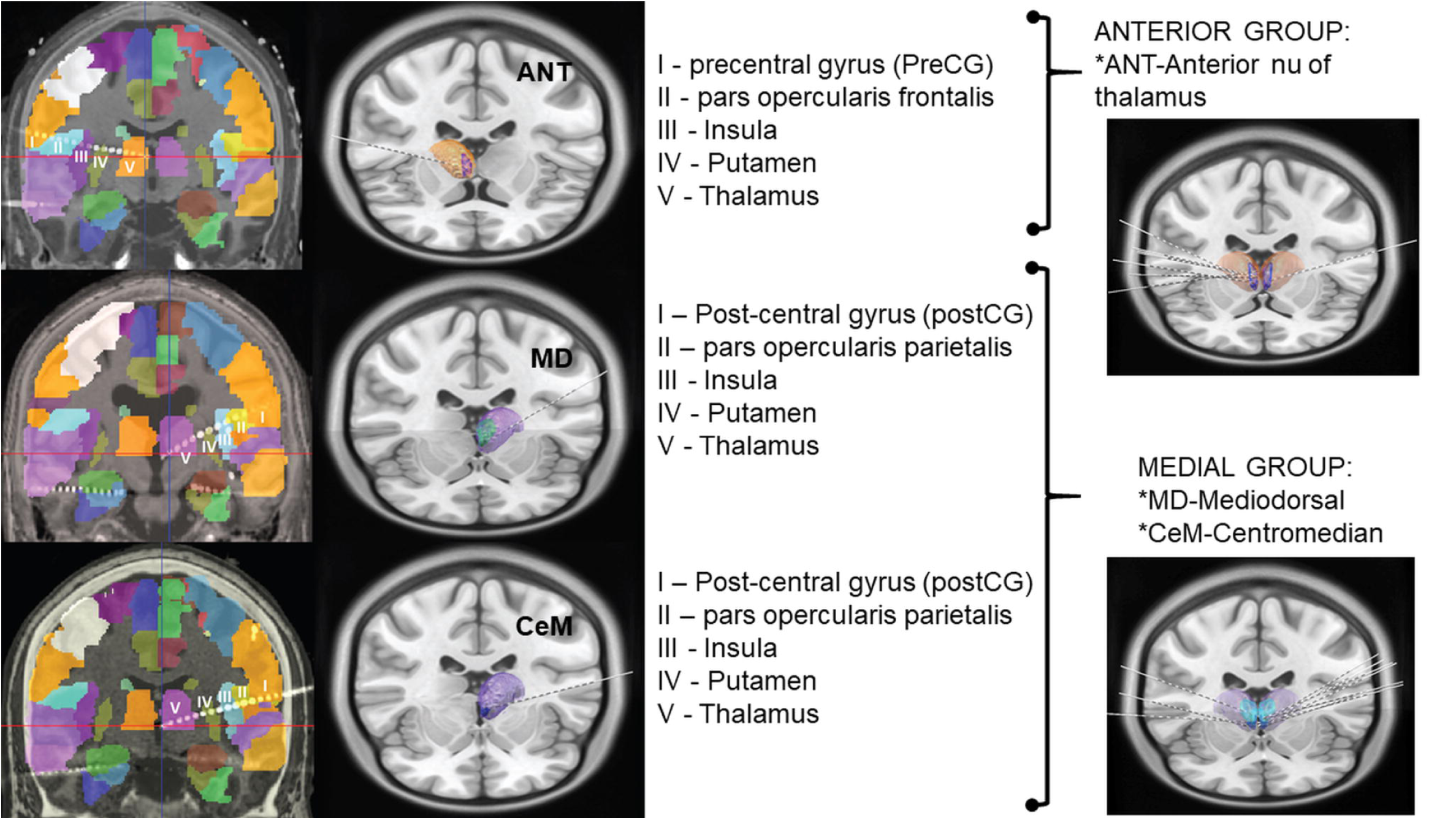
Implant registration accuracy: LeadDBS was used to reconstruct target location of the thalamic electrodes, while the trajectories were reconstructed and the cortical contacts were identified using iElectrodes. (A) ANT implantation followed the trajectory of precentral gyrus, pars opercularis frantalis, insula, putamen and thalamus, (B) medial group implantation followed similar trajectory except that the entry point was postcentral gyrus. (C&D) represent the group implants showing the trajectories of all ANT and medial group implants respectively.

Once the patients were implanted, the SEEG data would be recorded continuously. The thalamic SEEG signals in all patients were interpretable and comparable to the cortical channels. However, the local field potentials (LFPs) of the thalamus were of a lower amplitude compared to the cortical LFPs (Fig 6). The electrophysiological data thus obtained has been analyzed and published previously addressing key clinical questions^26,27,29,41^.

**Figure 6:**
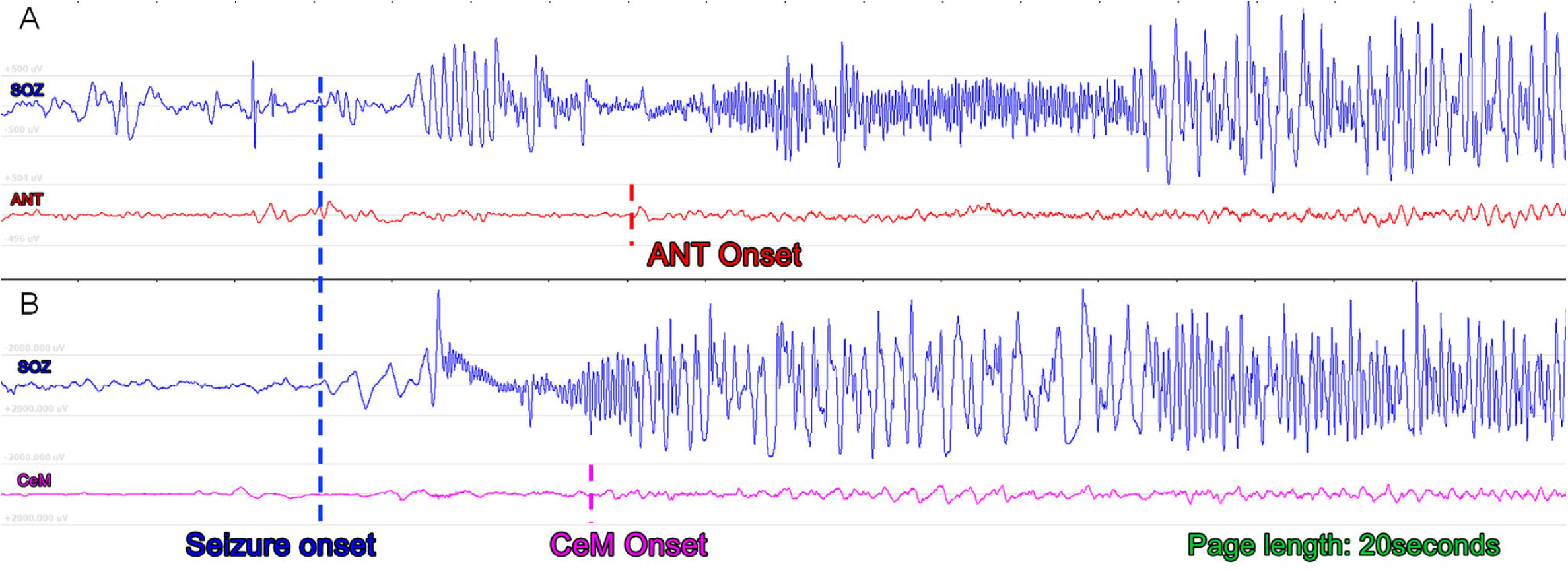
Seizure recording from the thalamus: (A) and (B) show seizures recorded in the thalamus. The identification of the unequivocal electrographic onset (UEO) was done by the clinician in the seizure onset channels (SOZ). Subsequent review of the thalamic channel showed that shortly after the seizure was noted in the SOZ, there was an evolving ictal rhythm noted in the thalamic channels. This EEG activity change was of a lower amplitude compared to that of the cortical channels and had a different morphology. (A) shows evolving ictal rhythm in a thalamic channel recording from ANT during a focal seizure originating in the ipsilateral hippocampus while (B) shows similar ictal activity recorded from CeM nucleus during a focal seizure originating from ipsilateral hippocampus.

## Discussion

Stereotactic procedures targeting the thalamus date back to the mid-20^th^ century when Spiegel and Wycis reported on thalamotomy for several psychiatric indications^33^ and they were also the first to record seizure activity from this location^38^. Since then, numerous thalamic stereotactic procedures have been performed for a wide variety of indications. By far, however, the most significant experience in thalamic stereotaxy has been implantation of deep brain stimulation (DBS) electrodes in the ventral intermediate thalamus (VIM) for the management of essential tremor. The published complication rates of these DBS procedures mirror SEEG overall, with a rate of 1-1.3% ^8,11,14^. McGovern et al in one of the pivotal studies has reported an overall hemorrhage rate of 19.1%^23^. while in our study, we had 20.8% hemorrhage rate, all of which were at the entry site of the electrodes. Contrary to the common apprehension that thalamic implant is associated with a high bleed rate, in our study we noted no thalamic bleeds. In the pivotal trial for ANT DBS, 4.5% of the 110 patients had incidental asymptomatic intracerebral hemorrhage but it is unclear if these bleeds were around the entry site of the DBS electrode or if they were thalamic. ^13,34^. Overall, SEEG requires an increased number of brain penetrations in comparison to DBS; however, the electrodes are smaller, and the procedure is very well-tolerated, with a reported hemorrhage rate of 1-4% ^8,11,14,24^. The higher complication rate reported in the study may be due to the difference in reporting. Prior SEEG studies have reported hemorrhage rate based on evaluating post-implant CT brain while in the present study, post-explant CT was used to estimate complications similar to McGovern et al. An electrophysiological sampling of the thalamus during SEEG investigation has been reported but complications were not studied in detail^1,12,15,30,31^.

Although the thalamus is successfully and safely targeted in DBS, SEEG requires quite a different trajectory planning and implantation techniques. In contrast to DBS, where there is considerable flexibility in the entry point, SEEG entries, .and target, as well as the structures along its path, are frequently constrained by MRI abnormalities, MEG or PET findings, intervening sulci, and vasculature. Also, thalamic DBS affords the opportunity in many cases to fully visualize the cortex if the dura is opened fully, whereas SEEG electrodes are placed via a small craniostomy in which direct visualization is not possible. Furthermore, a rigid cannula passed either to target or just shy of the target to guide placement of DBS electrodes, whereas SEEG electrodes are placed without a cannula, and are more prone to deviations. Finally, SEEG surgical planning requires achieving adequate coverage of the putative epileptogenic areas with a limited number of electrodes. In suspected TLE, anatomical sampling with SEEG often includes extra-temporal structures, including the insular operculum, orbitofrontal, parietal, and cingulate regions to rule of TLE mimics^2,20^. Numerous stereotactic techniques have been described for depth electrode placement including frame-based, frameless, and robotic methods each with their relative advantages and disadvantages. Many surgeons have recently gravitated towards robotic techniques due to the precision and speed offered. We do not endorse any particular technique, in fact, we believe thalamic depth electrode implantation is likely safe with any modern stereotactic system that has an accuracy error of approximately 1mm or less. We maintain that safety is more a factor of careful trajectory planning taking care to avoid cortical, sulcal, or deep vasculature; accurate image registration; and cautious surgical technique avoiding common complications such as drill skiving, drill plunging, and human measurement errors. These are fundamentals critical for the safety of all stereotactic procedures. Our study only utilized a single technique and therefore cannot directly address safety or accuracy differences in the various stereotactic methods.

Despite these myriad technical constraints, here we present results demonstrating that it is safe to extend clinically indicated trajectories, specifically those through the frontal operculum or insula, for accurate targeting of the limbic thalamus. The overall complication rates are low and comparable to electrodes placed in any other location. With these results, we propose that if patients are fully informed of the risks involved, there are significant benefits of obtaining robust signals from thalamic nuclei involved in seizure networks which may help guide future therapies ^26,27,29^. Some of the early studies by our group have shown (1) the periictal electrophysiological changes occurring in the thalamus during the seizures, (2) the cortical responsiveness to thalamic stimulation and (3) temporal predictive model to determine ictal and interictal thalamic states in TLE. In concordance with the growing evidence from various centers around the world, there is a possibility to envisage a more patient-oriented closed-loop DBS system. Systematic evaluation of complex thalamocortical interactions will eventually help in the development of such neuromodulation interventions in patients with drug-resistant epilepsies. Current thalamic DBS strategies are based mostly on a ‘one-size-fits-all’ model without a knowledge of the thalamocortical interactions specific to that patient. Estimating patient-specific inherent thalamocortical frequency interactions can help in tailoring the stimulation parameters and developing DBS systems to optimize clinical response which could significantly improve their clinical outcomes.

### Limitations

From our collective experience, we highlight some of the challenges and future perspectives about thalamic sampling during the SEEG investigation. First, target selection was performed directly on a 3T standard T1-weighted gadolinium-enhanced MRI sequences, which is challenging as both the ANT and MLT are not well visualized. While used as an external visual reference, the Morel’s thalamic atlas overlay or its coordinate system^21^ are not integrated into the robotic navigation systems to date, nor were specialized MR sequences to visualize landmarks such as the mammillothalamic tract on the FGATIR sequence. With experience and with potential incorporation of deformable atlases, we anticipate our targeting process will become more precise. Second, although no hemorrhagic or focal neurologic complications were noted, detailed neuropsychiatric examinations were not performed to assess whether the routine placement of electrodes produces damage that results in cognitive decline^42^. The randomized trial of ANT DBS did not find any cognitive decline with the placement of the ANT DBS. However, the transventricular trajectory utilized in that study is different than the lateral trajectory used in this study, making direct comparison difficult. Variable neuropsychological changes have been reported following ventral intermediate DBS that, when present, are thought to be primarily stimulation related and not a lesion effect^22,45^.

## Conclusions

The therapeutic potentials and prognostic role of the thalamus in focal epilepsy are well established in preclinical and clinical imaging studies. However, the lack of electrophysiological studies limits our knowledge of its involvement and may potentially hinder the development of therapies. Using robot-assisted SEEG, we demonstrate the safety of electrophysiological sampling from various thalamic nuclei for research recordings, presenting a technique that avoids implanting additional depth electrodes or comprising clinical care. We state with utmost caution that these results should not be taken for a safety blanket, but instead safe deep brain structure implantation should be judiciously performed through the development of meticulous anatomical target strategies and robotic planning to avoid untoward complications during the SEEG procedure.

## ABBREVIATIONS

SEEG: Stereoelectroencephalography
ANT: Anterior nucleus of thalamus
MED: Medial group of thalamic nuclei
MD: Mediodoral nucleus of thalamus
CeM: Centromedian nucleus of thalamus
MI: Massa-intermedia
ED: Euclidean Distance
TLE: temporal lobe epilepsy
EZ: Epileptogenic Zone

## Acknowledgments

The authors would like to acknowledge the contribution of patients and family members without whom this study would be incomplete. Ms. Jennifer Mahaffey and Ms. Cynthia Stover, Department of Neurology office, UAB, have extensively coordinated and organized the initial administrative components of the study. We would also like to recognize the continued enthusiasm and support from the EEG technicians at the UAB Epilepsy Center. SP and GC would like to acknowledge support from NIH (1RF1MH117155-01), and SP and AI would like to acknowledge support from NSF-EPSCoR (NSF RII□2FEC OIA1632891).

